# Macromolecular Crowding Promotes Reentrant Liquid-Liquid Phase Separation of Human Serum Transferrin and Prevents Surface-Induced Fibrillation

**DOI:** 10.1101/2023.06.21.545847

**Authors:** Chinmaya Kumar Patel, Chanchal Rani, Rajesh Kumar, Tushar Kanti Mukherjee

## Abstract

Protein aggregation and inactivation upon surface immobilization are major limiting factors for analytical applications in biotechnology related fields. Protein immobilization on solid surfaces often requires multi-step surface passivation which is time consuming and inefficient. Herein, we have discovered that biomolecular condensates of biologically active human serum transferrin (Tf) can effectively prevent surface-induced fibrillation and preserve the native-like conformation of phase separated Tf over a period of 30-days. It has been observed that macromolecular crowding promotes homotypic liquid-liquid phase separation (LLPS) of Tf through enthalpically driven multivalent hydrophobic interactions possibly via the involvement of its low complexity domain (residue 3–20) containing hydrophobic amino acids. The present LLPS of Tf is a rare example of salt-mediated reentrant phase separation in a broad range of salt concentrations (0–3 M) solely via the involvement of hydrophobic interactions. Notably, no liquid-to-solid-like phase transition has been observed over a period of 30-days, suggesting the intact conformational integrity of phase separated Tf as revealed from single droplet Raman, circular dichroism, and Fourier transform infrared spectroscopy measurements. More importantly, we discovered that the phase separated condensates of Tf completely inhibit the surface-induced fibrillation of Tf, illustrating the protective role of these liquid-like condensates against denaturation and aggregation of biomolecules. The cell mimicking aqueous compartments of biomolecular condensates with a substantial amount of interfacial water preserve the structure and functionality of biomolecules. Our present study highlights an important functional aspect of biologically active protein condensates and may have wide-ranging implications in cell physiology and biotechnological applications.

## Introduction

Membraneless cellular organelles formed via liquid-liquid phase separation (LLPS) of biomolecules play critical cellular functions via spatial and temporal organization of intracellular components. Various intracellular membraneless organelles in the cytoplasm and nucleus are formed via LLPS of proteins and/or nucleic acids.^1–3^ They are directly or indirectly involved in many cellular processes^4, 5^ including chromatin reorganization,^6^ noise buffering,^7^ sensing,^8^ signalling,^9, 10^ RNA storage/degradation,^11, 12^ T-cell activation,^13^ and ribosome biogenesis.^14^ Misregulation of these processes has been associated with serious pathological conditions.^15, 16^

Over the past decade, considerable efforts have been made to understand the fundamental mechanism behind the LLPS of various proteins, particularly intrinsically disordered proteins/regions (IDPs/IDRs) containing low-complexity domains/regions (LCDs/LCRs) in their polypeptide sequence.^17–35^ LLPS of biomolecules is a spontaneous and thermodynamically favorable liquid-liquid demixing phenomenon where the enthalpy change associated with the multivalent attractive intermolecular interactions, and the increase in entropy associated with the release of water molecules from the surfaces of biomolecules compensate the entropy loss due to the association of biomolecules. Due to this enthalpic and entropic contributions, the LLPS of biomolecules is highly sensitive on various internal and external factors including composition,^33^ concentration,^27^ temperature,^31^ pressure,^36^ pH,^26^ ionic strength,^37^ and crowding.^21, 22^ The phase separated biomolecule-rich phase contains highly dynamic liquid-like membraneless condensates which are stabilized by various multivalent intermolecular forces including electrostatic, hydrophobic, hydrogen bonding, dipole-dipole, π-π, and/or cation-π interactions.^18, 24, 33^ Recent studies have demonstrated that these membraneless condensates act as dynamic intermediates during the formation of amyloid aggregates of various disease-associated proteins or polypeptides.^19–22, 27, 28, 38–43^ Earlier, Maji and coworkers quantified the conformational heterogeneity during liquid-to-solid phase transition of α-Synuclein and shown that the interior of phase separated droplets is heterogeneous with the presence of monomers, oligomers, and fibrils.^21, 22^ More importantly, it has been observed that the phase-separated droplets act as nucleation scaffolds for protein aggregation and liquid-to-solid phase transition. Recently, Vendruscolo and coworkers studied the aggregation kinetics of α-synuclein within liquid condensates and shown that α-synuclein can undergo spontaneous homogeneous primary nucleation and fast aggregate-dependent proliferation within condensates at physiological pH.^43^ Similarly, using a family of chimeric peptides (ACC_1–13_K_n_), Winter and coworkers demonstrated that the early fibrils originate at the droplet/bulk interface within single liquid droplets in the ATP-triggered amyloidogenic pathway.^38^ While the interior of these biomolecular condensates are highly dynamic and heterogeneous in nature, their solid-like fibrillar aggregates have relatively defined and homogeneous structural composition. Although much is understood about the LLPS of disease-associated proteins and their aggregation into solid-like fibrillar aggregates, little has been uncovered about the role of these condensates on the activity and properties of functional native proteins. Very recently, it has been discovered that functional globular proteins lacking any IDRs/LCDs can also undergo LLPS in the presence of inert macromolecular crowders via intermolecular protein-protein interactions.^44, 45^ The nature and extent of these weak protein-protein interactions depend on the overall structure and amino acid composition of individual proteins. More importantly, our group has recently demonstrated macromolecular crowding-induced biomolecular condensate formation of two structurally robust functional enzymes, namely horseradish peroxidase (HRP) and glucose oxidase (GOx), which remarkably enhances the enzymatic activity and selectivity at physiological conditions.^46^ These intriguing findings of homotypic LLPS of functional proteins open up a new avenue for the exploration of the role of these membraneless condensates to control the activity of various functional proteins. Apart from their role in aggregation pathway, various synthetic and biomolecular condensates are also known as efficient scaffolds for the stabilization and protection of sequestered biomolecules against temperature, pH, protease digestion, and aging.^47–49^ In the present study, we have explored the phase behavior of an important biologically active protein namely, human serum transferrin (Tf) in the absence and presence of inert macromolecular crowders under physiological conditions.

Tf is a bilobal iron transport glycoprotein with a molecular weight of 80 kDa, which circulates in blood at a concentration of ∼35 µM and has a half-life of 7–8 days.^50–53^ It is synthesized in the liver and subsequently secreted into the plasma. Tf has a single polypeptide chain with 698 amino acid residues and contains two homologous lobes (N-lobe and C-lobe) of an equal weight of 40 kDa connected by a short flexible hinge (Scheme 1a).^54–56^ Each lobe consists of two sub domains (N1, N2 and C1, C2) with an octahedral Fe^3+^ binding site. Each Fe^3+^ ion is coordinated with four highly conserved residues, namely an aspartic acid, two tyrosine, and a histidine along with a bidentate carbonate anion.^51^ The two lobes are in an open conformation in the absence of bound Fe^3+^ in apotransferrin (apo-Tf). The release of bound iron is triggered by a drop in pH as a consequence of protonation of the carbonate and other amino acids in the interdomain binding cleft. Abnormal and uncontrolled aggregation of Tf is linked with irregular iron distribution and iron overload, which may have serious pathological conditions including neurogenerative diseases.^50, 51^ The polypeptide chain of holo-Tf has a LCD in the range of residue number 3–20, as revealed from the SMART analysis (Scheme 1b).^57^ This 18-residue long LCD falls in the N-lobe of the holo-Tf and mostly consists of hydrophobic amino acids with the composition of six leucine, four alanine, four valine, two glycine, and two cysteine residues. On the other hand, IUPred2 analysis reveals mostly ordered conformation of holo-Tf with two very short disorder segments (residues 555–558, and 561–570) in the C-lobe (Scheme 1c).^58^

**Scheme 1.**
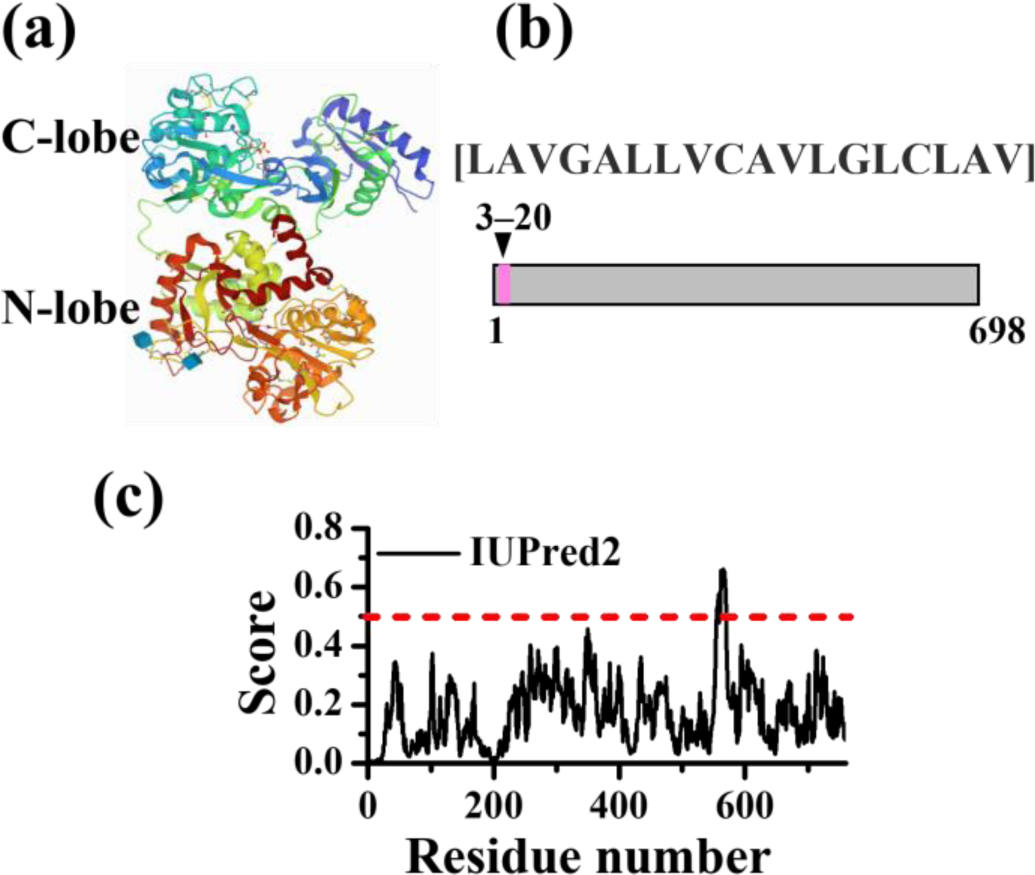
(a) Crystal Structure of Tf (PDB entry 3QYT) Obtained from the Protein Data Bank. (b) Analysis of the Primary Sequence of Human Serum Transferrin Using SMART Algorithm to Predict the LCDs, and (c) IUPred2 Analysis to Predict the Disorder Tendency.

These two segments are mainly composed with polar (glutamine, threonine, asparagine), charged (lysine, aspartic acid) as well as hydrophobic (proline, glycine, valine, tryptophan) amino acid residues with random sequences. The role of the distinct LCD with hydrophobic amino acid residues on the LLPS of Tf has not been explored yet. We envisaged that the presence of a distinct LCD might facilitates Tf to undergo phase separation under optimized physiological conditions. However, it should be noted that the presence of IDRs and/or LCDs is not a mandatory prerequisite for biomolecules to undergo LLPS.^44–46^ To better understand the underlying factors that govern the LLPS and functional aspects of membraneless condensates of biologically active proteins, we have undertaken the present study. In particular, we want to know whether Tf undergoes LLPS and subsequent aggregation via liquid-like condensate formation within the cellular lifetime of Tf under physiological conditions.

## Experimental Section

### Materials

Human serum transferrin (Tf), human apo-transferrin (apo-Tf), polyethylene glycol 8000 (PEG 8000), Ficoll 400, bovine serum albumin (BSA), dextran 70, sodium thiocyanate (NaSCN), 1,6-hexanediol, rhodamine B isothiocyanate (RBITC), fluorescein-5-isothiocyanate (FITC), proteinase K, thioflavin T (Th-T), hellmanex III and the Pur-A-Lyzert dialysis kit (molecular weight cutoff 3.5 kDa) were purchased from Sigma-Aldrich. Sodium dihydrogen phosphate monohydrate (NaH_2_PO_4_.H_2_O), di-sodium hydrogen phosphate heptahydrate (Na_2_HPO_4_.7H_2_O), sodium acetate trihydrate (CH_3_COONa.3H_2_O), acetic acid (CH_3_COOH), sodium bicarbonate (NaHCO_3_), sodium carbonate anhydrous (Na_2_CO_3_), sodium chloride (NaCl), sodium azide (Na_3_N) and methanol (MeOH) were purchased from Merck. All the chemicals were used without any further purification. Eco Testr pH1-pH meter was used to adjust the final pH (± 0.1) of all the buffer solutions. Milli-Q water was obtained from a Millipore water purifier system (Milli-Q integral).

### Characterization Techniques

#### Confocal Laser Scanning Microscopy (CLSM)

The confocal images were captured using an inverted confocal microscope, Olympus fluoView (model FV1200MPE, IX-83) through an oil immersion objective (100 × 1.4 NA). The samples were excited using two different diode lasers (488 and 559 nm) by using appropriate dichroic and emission filters in the optical path. Confocal images were captured for both dried and liquid samples. For surface-dried experiments, samples (∼20 µL) were drop-cast onto a cleaned coverslips and dried overnight inside a desiccator. For liquid phase experiments, 20 μL aliquot of the sample solution was drop-cast onto a cleaned glass slide and sandwiched with a Blue Star coverslip. Finally, the sides of the coverslips were sealed with commercially available nail paint, and then images were captured. The prepared samples were placed in a desiccator under vacuum and left overnight for drying.

#### Field Emission Scanning Electron Microscopy (FESEM)

The FESEM images were obtained using a field-emission scanning electron microscope, supra 55 Zeiss. For FESEM measurements, the samples were drop-cast on a cleaned coverslip and were dried overnight in a desiccator. Subsequently, the dried samples were coated with gold and imaged under the microscope.

#### Fourier-Transform Infrared (FTIR) Spectroscopy

FTIR measurements were performed to determine the secondary structure of the proteins using a Bruker spectrometer (Tensor-27). 10 µL of each liquid sample (50 µM Tf in the presence/absence of different crowders and in different pH buffers) was used for FTIR measurement. The spectra were recorded in the range of 4000–400 cm^−1^. Fourier self-deconvolution (FSD) method was used to deconvolute the spectra corresponding to the wavenumber 1700−1600 cm^−1^.^59^ The Lorentzian curve fitting was done to fit the individual spectrum using origin 8.1 software.

#### Circular Dichroism (CD) Spectroscopy

CD spectra were recorded on a JASCO J-815 CD spectropolarimeter using a quartz cell of 1 mm path length with a scan range of 190-260 nm. Spectra were recorded with a slit width of 1 mm and speed of 50 nm/min. These spectra were plotted using origin 8.1 software.

#### Raman Spectroscopy

Raman spectra were recorded using Horiba-Jobin Yvon micro-Raman spectrometer equipped with a 532 nm excitation laser and CCD detector. Samples (20 µL aliquots) of Tf fibril (50 µM Tf in phosphate buffer) and Tf droplet (50 µM Tf + 10% PEG in phosphate buffer) were deposited on the cleaned coverslips and kept inside an incubation chamber at 37 °C for 1-to 14-days. A 532 nm excitation laser light with 40 mW (100%) laser power was utilized to excite the samples. The laser was focused using a 50× objective lens. The spot size of laser beam in the Raman spectrometer is ∼2 µm and the diffraction grating of 600 lines/mm were used to disperse the scattering light. Data were collected in the backscattering mode with flat correction using inbuilt method in Raman spectrometer software.

The obtained experimental spectra were baseline corrected and normalized with respect to phenylalanine ring breathing band at 1002 cm^−1^ using origin 8.1 software.

## Results and Discussion

### Macromolecular Crowding Promotes LLPS of Tf

Recent studies have demonstrated that inert synthetic/protein crowders can effectively modulate the intermolecular protein-protein interactions and facilitate LLPS via lowering the critical concentration of protein required for LLPS.^19–22, 32, 44–46^ In the present study, we have investigated the phase behavior of Tf in the absence and presence of different inert synthetic crowders such as polyethylene glycol (PEG 8000), dextran 70, and Ficoll 400 along with inert protein crowder, namely bovine serum albumin (BSA) to mimic the intracellular crowding. Phase behavior studies were performed in liquid chamber under the confocal laser scanning microscope (CLSM) using fluorescein-5-isothiocyanate (FITC)-labelled Tf in the concentration range of 1–50 µM (Figure 1a). In the absence of PEG 8000, holo-Tf shows no characteristic features in the phase contrast as well as in fluorescence images (Figure 1b). However, under the same experimental conditions, distinct spherical assemblies are observed in the presence of 10% PEG 8000 (Figure 1b). Notably, the uniform fluorescence signals appear exclusively from the interior of these assemblies, signifying the presence of FITC-labelled Tf. These spherical assemblies exhibit characteristic liquid-like properties such as spontaneous fusion, surface wetting, and dripping (Figures 1c and S1), suggesting that these assemblies are liquid-like coacervates of Tf. In order to know whether these coacervates are formed due to any specific interactions of Tf with PEG 8000 or it is a generic phenomenon, we performed the phase separation experiments under similar experimental conditions in the presence of other polymeric and protein crowders, namely 12.5% Ficoll 400, 10% dextran 70, and 20 mg/mL BSA. Notably, Tf form similar spherical green emitting coacervates in the presence of Ficoll, dextran, and BSA (Figures 1d), signifying a generic crowding effect. Similar crowding-induced phase separation and liquid-like condensate formation has also been observed with apo-Tf under the same experimental conditions (Figure S2). The liquid phase aging of 1 µM Tf solution in the presence of 10% PEG reveals increase in the mean size of Tf droplets over a period of 14-days due to the spontaneous coalescence/fusion events (Figures S3 and S4).

**Figure 1.**
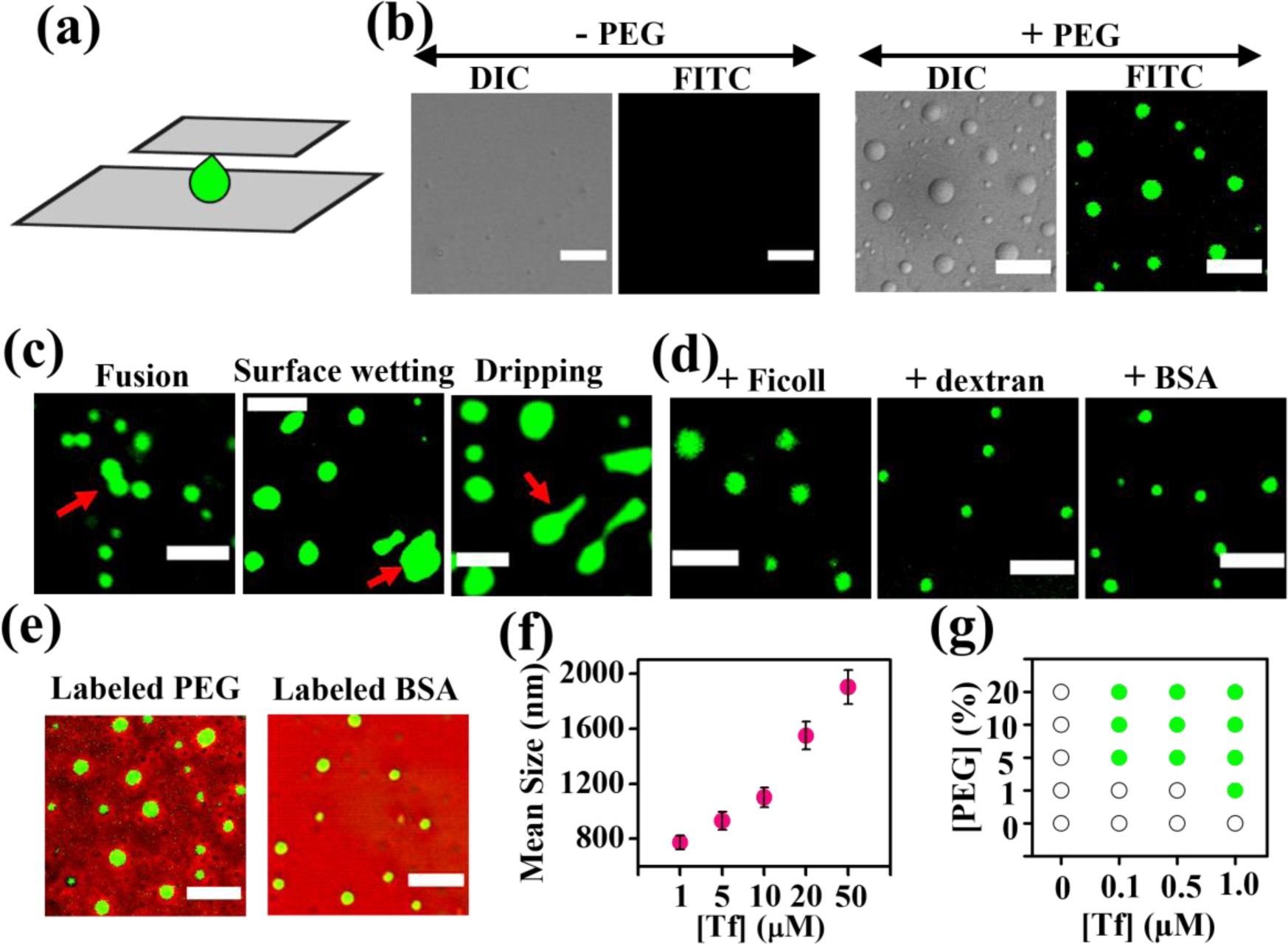
(a) Schematic illustration showing the liquid compartment for confocal imaging. (b) Confocal (DIC and fluorescence) images of FITC-labelled 1 µM Tf in the absence and presence of 10% PEG. (c) Confocal images of Tf droplets showing fusion, surface wetting and dripping phenomenon. The red arrows highlight these phenomena. Confocal images showing droplet formation of FITC-labelled 1 µM Tf in the presence of (d) unlabeled 12.5% Ficoll 400, 10% dextran 70, and 20 mg/mL BSA and (e) RBITC-labeled 10% PEG and 20 mg/mL BSA at 37 ℃. (f) Changes in the mean size of Tf droplets estimated from the confocal images as a function of Tf concentrations in the presence of 10% PEG after 1-day of incubation. The data points represent the mean ± s.e.m. for three independent measurements. (g) Phase diagram of Tf as a function of PEG concentrations at 37 ℃. The scale bars correspond to 5 µm.

The formation mechanism of biomolecular condensates can be either homotypic (only proteins are involved) or heterotypic (with foreign crowders) in nature. To know whether the present condensates are homotypic or heterotypic in nature, we performed phase separation assay under confocal microscope with FITC-labeled Tf in the presence of rhodamine B isothiocyanate (RBITC)-labeled PEG (mPEG-NH_2_, MW 5000) and BSA. Interestingly, the red background emissions from RBITC-labeled PEG and BSA are totally excluded from the distinct green emissions from FTIC-labeled Tf droplets (Figure 1e). These observations substantiate that the present condensates are formed via homotypic LLPS of Tf in the presence of inert crowders which are entirely excluded from the condense phase. The mean size of these condensates increases from a value of 774 ± 50 to 1903 ± 123 nm upon increase in the concentrations of Tf from 1 to 50 µM in the presence of 10% PEG (Figures 1f and S5). This observation indicates the dominant role of protein-protein interactions at higher concentrations of Tf. The phase diagram of Tf as a function of PEG concentrations reveals that the critical concentration of Tf required for phase separation decreases with increase in the macromolecular crowding (Figure 1g). This can be explained by considering enhanced protein-protein interactions at lower concentration of Tf due to the excluded volume effect of macromolecular crowders. Taken together, our findings indicate that Tf undergo spontaneous homotypic LLPS via formation of liquid-condensates upon macromolecular crowding by means of intermolecular protein-protein interactions.

### Effect of Temperature and pH on LLPS of Tf

Temperature and pH of the medium have enormous impact on the structure and function of proteins and both these parameters directly influence the LLPS of various proteins.^26, 44–46^ Therefore, we investigated the influence of temperature and pH of the medium on the extent and feasibility of LLPS of Tf. Depending on the molecular driving forces, proteins can either exhibit upper critical solution temperature (UCST)^17^ or lower critical solution temperature (LCST)^19^ or both.^26^ To know the effect of temperature on the LLPS of Tf, we directly monitored the feasibility of LLPS of 1 µM Tf in the presence of 10% PEG 8000 as a function of temperature in the range of 4–50 ℃ using CLSM (Figure 2a). Our findings reveal that the biomolecular condensates of Tf are structurally stable in the temperature range between 4–37 ℃. The mean size of these condensates increases from 459 to 774 nm upon increasing the temperature from 4–37 ℃. Notably, these liquid-droplets are unstable at and beyond 50 ℃ (Figure 2a), indicating that the phase separation of Tf follows a UCST phase profile.

**Figure 2.**
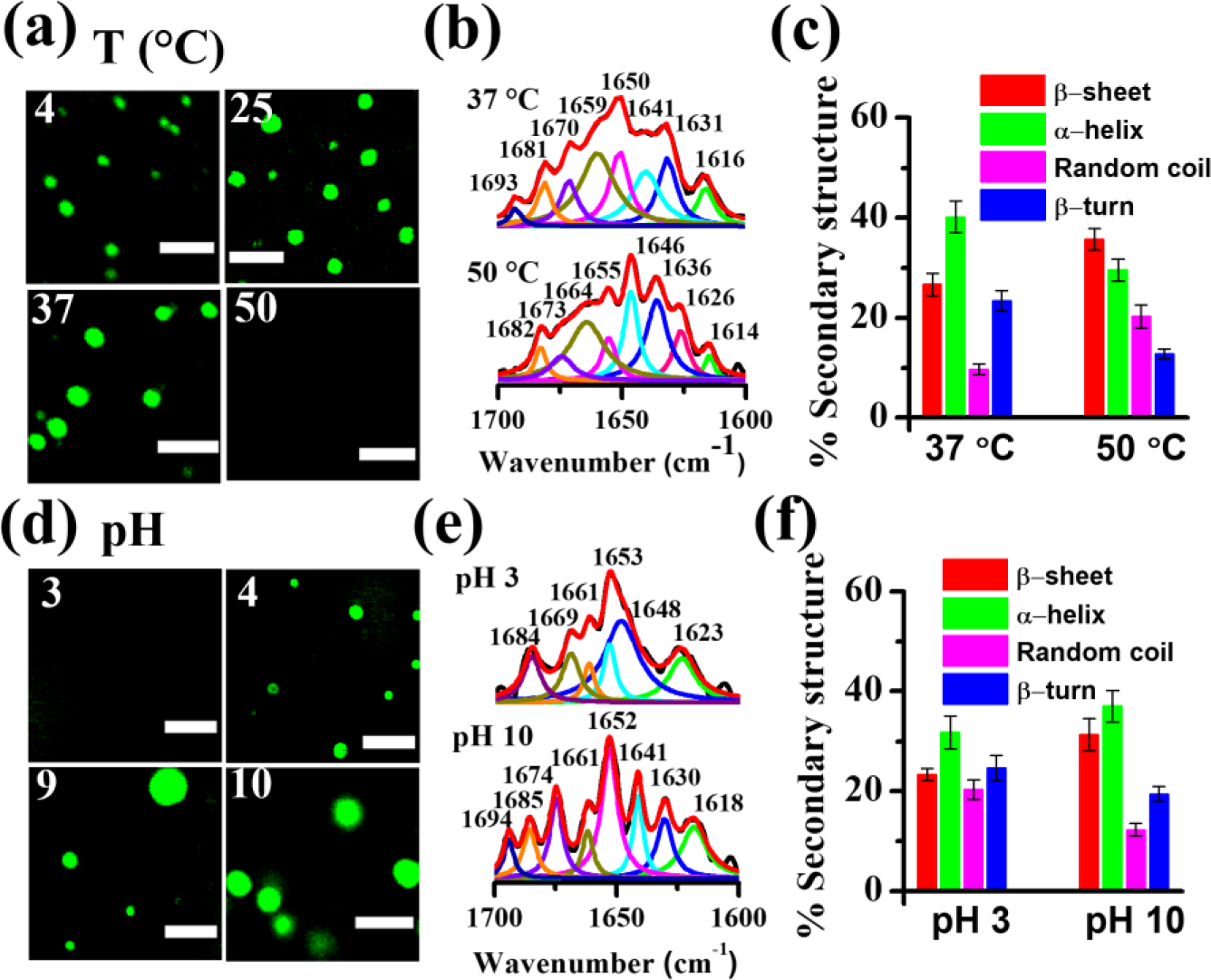
(a) Confocal images showing the effect of temperatures on the stability of FITC-labeled Tf droplets in the presence of 10% PEG. (b) Deconvoluted FTIR spectra of 50 µM Tf in the presence of 10% PEG at 37 and 50 °C. (c) Determination of the percentage of secondary structures of Tf in the presence of 10% PEG at 37 and 50 °C from the deconvoluted FTIR spectra. The data points represent the mean ± s.e.m. for three independent measurements. (d) Confocal images showing the effect of pH on the stability of FITC-labeled Tf droplets in the presence of 10% PEG at 37 °C for 1-day. (e) Deconvoluted FTIR spectra of 50 µM Tf in the presence of 10% PEG at pH 3.0 and pH 10.0. (f) Determination of the percentage of secondary structures of Tf in the presence of 10% PEG at pH 3.0 and pH 10.0 from the deconvoluted FTIR spectra. The data points represent the mean ± s.e.m. for three independent measurements. The scale bars correspond to 5 μm.

Similar UCST phase profiles have been observed previously for several other proteins.^17, 26, 31, 44– 46^ The present UCST behavior of Tf suggests that the phase separation of Tf is mostly driven by enthalpically favorable multivalent intermolecular protein-protein interactions between Tf molecules. These soft interactions are mainly short-range and may include electrostatic, hydrophobic, π-π, cation-π, and/or hydrogen bonding.^18, 24, 33^ Importantly, the LLPS of biomolecules is energetically favourable only when the enthalpy gain from these weak multivalent attractive interactions overcomes the entropy of mixing as predicted previously by Flory–Huggins theory.^60, 61^ The observed inhibition of the LLPS of Tf at 50 ℃ might be due to the altered secondary structure, which ultimately modulates the protein-protein interactions. To examine the extent of alteration of the secondary structure of Tf as a function of temperature, we recorded FTIR spectra at 37 and 50 ℃ (Figure 2b). The deconvoluted FTIR spectra of Tf at 37 ℃ reveal the presence of a major peak at ∼ 1650 cm^−1^, which is a characteristic peak of the α-helical structure. In contrast, noticeable changes in the spectral signatures of Tf at 50 ℃ have been observed. Tf at 50 ℃ shows a major peak at ∼ 1646 cm^−1^, which is a characteristic peak of random coil structure (Figure 2b). Notably, a significant rise in the β-sheet and random coil contents at 50 ℃ suggests a prominent conformational alteration of Tf (Figure 2c), which prevents the LLPS of Tf via weakening of the intermolecular protein-protein interactions. In order to know the effect of pH on the LLPS of Tf, the solution pH was varied from 3.0–10.0. The CLSM images reveal that the droplets of Tf are structurally stable in the pH range of 4.0–10.0, however, phase separation is completely inhibited at a pH value of 3.0 (Figure 2d). It is known that pH has a profound influence on the secondary structure of proteins and can modulate the feasibility of LLPS. To know the extent of modulation of the secondary structure of Tf at lower acidic pH of 3.0, we compared the deconvoluted FTIR spectra of Tf at pH 10.0 and 3.0 (Figure 2e). A noticeable rise in the random coil and *β*-turn contents along with a slight decrease in the α-helix content is clearly evident at pH 3 relative to that in pH 10.0 (Figure 2f). These observations are further supported from the CD measurements (Figure S6). CD spectral changes reveal appreciable changes in the secondary structures of Tf at pH 3.0 relative to that in neutral or basic pH. Previous studies have shown that in acidic medium (≤ pH 3.5) Tf loses its bound irons via a conformational change from close to an open form.^51–55^ It should be noted that, the isoelectric point of holo-Tf is ∼5.2,^53^ and hence, it bears net negative and positive surface charges at physiological pH of 7.4 and acidic pH of 4.0, respectively. Nevertheless, we observed that Tf undergoes spontaneous LLPS at both the pH values of 7.4 and 4.0, suggesting that the overall surface charge has negligible influence on the phase separation of Tf. Taken together, our findings reveal that the native protein structures play an important role in driving the LLPS via protein-protein interactions.

### Nature of Intermolecular Protein-Protein Interactions

To understand the mechanism of LLPS of Tf, it is important to decipher the nature of various intermolecular protein-protein interactions. Control experiments were performed by varying the concentrations of salts and aliphatic alcohols which are known to modulate the protein-protein interactions and perturb the LLPS of different proteins.^17, 18, 44–46^ Chaotropic salt sodium thiocyanate (NaSCN), and aliphatic alcohol 1,6-hexanediol are known to disrupt the hydrophobic protein-protein interactions. We visualized the feasibility of LLPS of 1 µM Tf in the presence of 10% PEG using CLSM by varying the concentrations of NaSCN from 0.1–2.0 M (Figures 3a,b and S7). Droplet formation is feasible up to 1.0 M of NaSCN; however, complete inhibition of LLPS of Tf has been observed at and beyond 2 M of NaSCN. This finding suggests the possible role of short-range hydrophobic protein-protein interactions behind the observed LLPS of Tf. To further substantiate the role of hydrophobic interactions, we varied the concentrations of 1,6-hexanediol. While the presence of 1,6-hexanediol up to a concentration of 3.0% has negligible effect on the phase separation of Tf, further increase in the concentration to 6.0% or above leads to a complete inhibition of the droplet formation (Figures 3a,b and S8).

**Figure 3.**
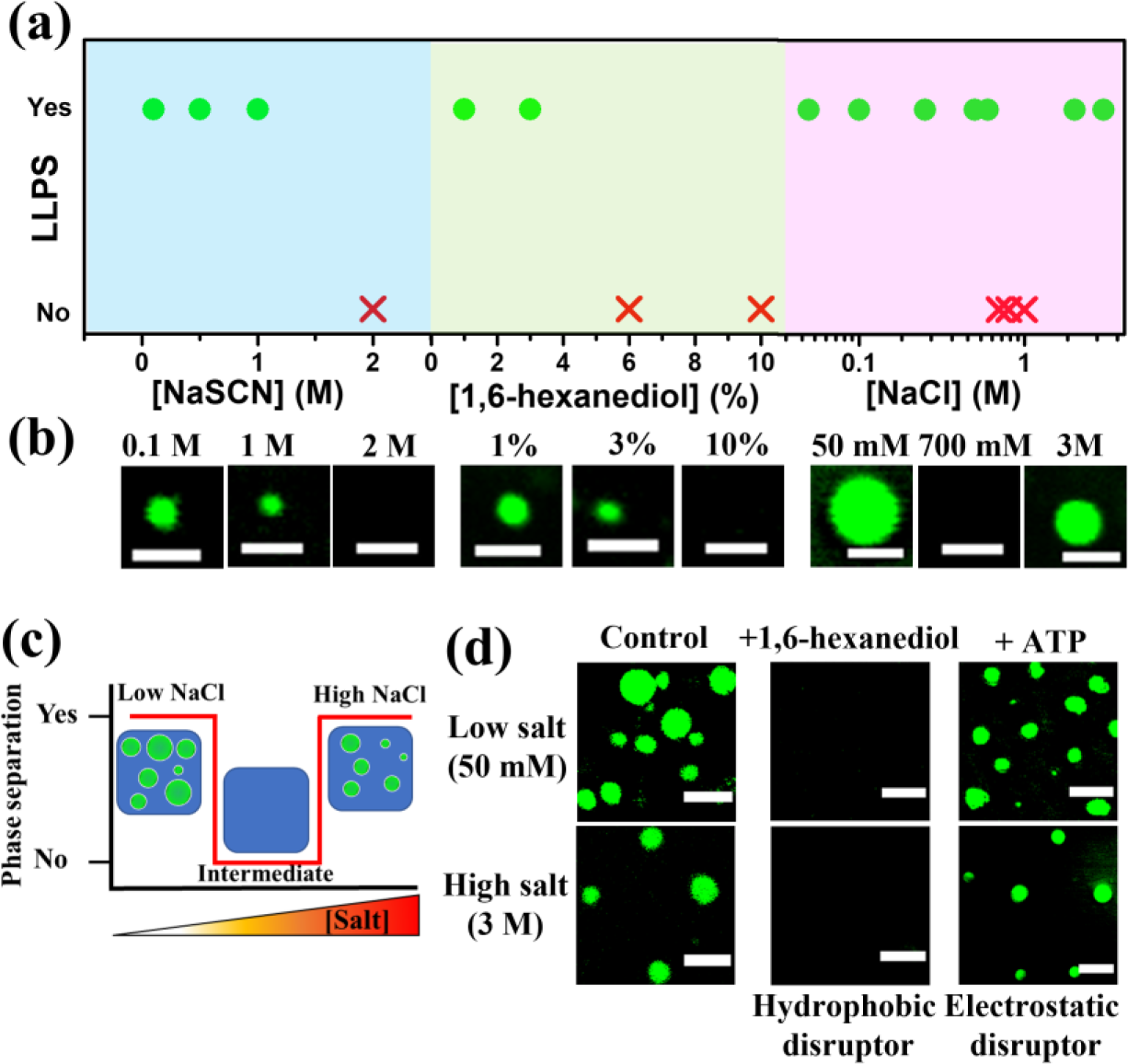
(a) Phase profile diagrams showing the LLPS propensity of 1 µM Tf in the presence 10% PEG as a function of NaSCN, 1,6-hexanediol, and NaCl concentrations at 37 °C upon 1-day of aging. (b) Representative confocal images of FITC-labelled 1 µM Tf droplets in the presence of different concentrations of NaSCN, 1,6-hexanediol, and NaCl concentrations incubated at 37 °C for 1-day. The scale bars correspond to 2 μm. (c) Schematic illustration of salt-mediated reentrant LLPS of Tf. (d) Confocal images of FITC-labelled Tf droplets in the low- and high-salt regimes in the absence and presence of 10% 1,6-hexanediol and 12.5 mM ATP. The scale bars correspond to 5 μm.

Furthermore, ammonium sulfate (NH_4_)_2_SO_4_ which is a kosmotropic salt favours the LLPS of Tf as revealed from the CLSM measurements (Figure S9). These findings validate the active role of hydrophobic protein-protein interactions behind the observed phase separation of Tf in the presence of macromolecular crowders. Next, to know the role of electrostatic interactions on the LLPS of Tf, we varied the concentrations of NaCl in the concentration range of 50 mM to 3.0 M (Figures 3a,b and S10). To our surprise, a biphasic reentrant phase separation behavior of Tf is observed as a function of NaCl concentrations. Tf forms phase separated droplets up to a NaCl concentrations of 600 mM and beyond that it exhibits a well-mixed homogeneous phase between 700 mM and 1.0 M of NaCl. Interestingly, Tf reenters the phase-separated regime at and beyond 2.0 M of NaCl. Although, this kind of reentrant phase behavior is very uncommon for homotypic LLPS of functional biomolecules at physiological conditions, similar salt-mediated reentrant phase separation has been reported recently by Knowles and coworkers for neurodegenerative disease associated proteins.^25^ The present biphasic LLPS of Tf is fundamentally different from that reported previously for RNA-mediated heterotypic reentrant phase behavior of protein-RNA condensates.^62, 63^ While the protein-RNA condensates were reported to be stable only in the presence of intermediate RNA concentrations, they remain as well-mixed homogeneous phase at both high and low RNA concentrations. Furthermore, reentrant phase separation of macromolecules has also been anticipated recently as a function of temperature,^64^ pH,^65^ and pressure.^66^ Here it is important to note that the molecular forces responsible for phase separation at high-salt regime may or may not be same with that at low-salt regime. For instance, Knowles and coworkers reported salt-dependent reentrant phase behavior of FUS and observed that both hydrophobic and electrostatic interactions drive the phase separation at low-salt concentrations, whereas the reentrant high-salt phase separation is primarily driven by hydrophobic interactions.^25^ To understand this unusual biphasic phase separation phenomenon (Figure 3c), we determined the molecular forces responsible for the phase separation of Tf in the low-salt and high-salt regimes by using hydrophobic (1,6-hexanediol) and electrostatic (ATP) disruptors (Figure 3d). For this, we have utilized Tf droplets formed in the presence of 50 mM and 3.0 M NaCl in 10% PEG. Notably, complete dissolution of Tf droplets has been observed upon addition of 10% 1,6-hexanediol both in 50 mM and 3 M NaCl, suggesting that hydrophobic interactions primarily drive the LLPS of Tf both in low-salt and high-salt regimes. In order to know whether electrostatic interaction has any role in the LLPS of Tf at low-salt and high-salt regimes, we utilized 12.5 mM ATP as electrostatic disruptor. Interestingly, addition of ATP has no effect on the feasibility of LLPS of Tf both in low-salt and high-salt regimes, suggesting negligible role of electrostatic interactions behind the LLPS of Tf. Therefore, these findings reveal that the homotypic reentrant phase separation of Tf in the presence of macromolecular crowders is primarily driven by hydrophobic protein-protein interactions. Here it is important to comment on the possible role of LCD of Tf on the observed LLPS. The amino acid composition and in particular the specific sequence of LCDs determine their ability to drive the LLPS of LCD containing proteins. Previously, LCD-mediated LLPS has been demonstrated for various IDPs including hnRNPA1,^17^ DEAD-box helicases Ddx4,^67^ and Laf-1.^35^ Our findings indicate that the LLPS of Tf is enthalpy driven and multivalent hydrophobic interactions are sole driving force both in low-salt and high-salt regimes. Indeed, careful scrutiny of the amino acid composition of the 18-residue long LCD in the N-lobe of Tf reveals the presence of mostly hydrophobic amino acids with the composition of six leucine, four alanine, four valine, two glycine, and two cysteine residues. This LCD of Tf may thus acts as an active interaction motif for the associative protein-protein interactions and enable multivalent hydrophobic contacts that drive the LLPS of Tf.

### Alteration of Secondary Structure of Phase Separated Tf

Upon establishing the molecular driving force behind the LLPS of Tf in the presence of inert crowders, we next examined the impact of the hydrophobic protein-protein interaction on the secondary structure of Tf after the LLPS. CD spectrum Tf shows a negative peak at ∼ 208 nm with a weak shoulder at ∼ 222 nm, indicating a α-helix rich secondary structure (Figure 4a). Notably, no spectral changes have been observed upon addition of 10% PEG even for a period of 14-days (Figure 4a).

**Figure 4.**
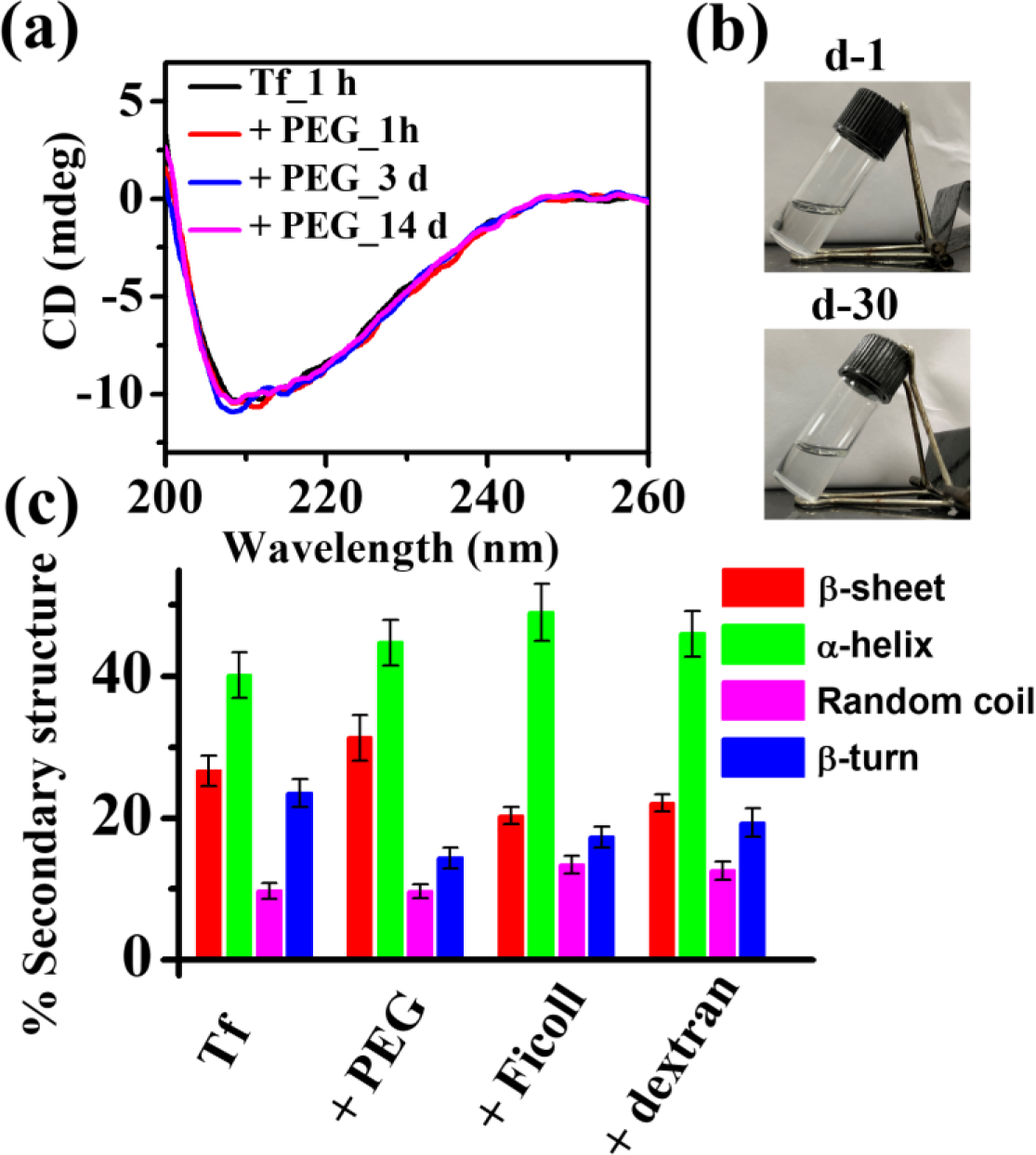
(a) CD spectra of 1 µM Tf in the absence and presence of 10% PEG over 14-days of solution phase aging. (b) The daylight photographs of 1 µM Tf solutions in the presence of 10% PEG aged for d-1 and d-30. (c) Plot showing the percentage of secondary structures of Tf in the absence and presence of various crowders estimated from the deconvoluted FTIR spectra. The data points represent the mean ± s.e.m. for three independent measurements.

Furthermore, no liquid-to-solid-like phase transition has been observed as revealed from CD spectrum and day-light photograph of a 30-days aged sample (Figure 4b). This suggests that the LLPS of Tf via hydrophobic protein-protein interactions has subtle influence on the secondary conformation of Tf in the presence of inert crowders. To further substantiate this argument, we recorded FTIR spectra of Tf in the presence of different crowders (Figure S11). It is evident that a major peak at 1652 nm has been observed irrespective of the nature of crowders, signifying that the overall α-helix rich structure of Tf remains unaltered upon LLPS in the presence of inert crowders (Figure 4c). Taken together, our findings reveal that the intermolecular hydrophobic interactions which drive the LLPS of Tf are soft in nature and do not perturb the overall secondary structure of Tf.

### Liquid Condensates Prevent Surface-Induced Fibrillation of Tf

Along with various intracellular functions, membraneless condensates are also linked with the abnormal protein aggregation/fibrillation in neurodegenerative diseases.^17, 18, 21, 27, 28, 32, 38^ Moreover, synthetic condensate as a scaffold can also acts as a protective shell to stabilize the sequestered biomolecules against various external stimuli.^49^ The spontaneous formation of liquid-like condensates of Tf via LLPS at physiological conditions suggests that these condensates might have important physiological functions. However, very less is known about the in vitro functional aspects of these membraneless condensates formed via homotypic LLPS of functional proteins. Herein, we investigated and compared the structural and morphological stabilities of individual bare Tf with the phase separated Tf droplets on a solid glass surface. The aqueous solution of 1 µM holo-Tf was prepared in pH 7.4 phosphate buffer at 37 ℃ in the absence and presence of 10% PEG and subsequently drop-casted on a glass surface for confocal measurements (Figure 5a). Samples were stored in a constant temperature incubation chamber for different time intervals (1–30 days). Figure 5b shows the DIC images of surface-deposited bare Tf in the absence of 10% PEG. It is evident that bare Tf undergoes aggregation with fibrillar morphology upon surface deposition (d-1). Notably, further aging of the drop-casted bare Tf sample for up to 14-days does not results in any significant change in the fibrillar morphology. Similar fibrillar aggregates have also been observed in FESEM measurements (Figure S12). In order to authenticate that these fibrillar aggregates are formed solely via the aggregation of Tf, we fluorescently labeled Tf molecules with green emissive FITC dye and imaged them under CLSM. Figure 5c displays the CLSM images of FTIC-labeled Tf on the glass surface upon d-1 of incubation.

**Figure 5.**
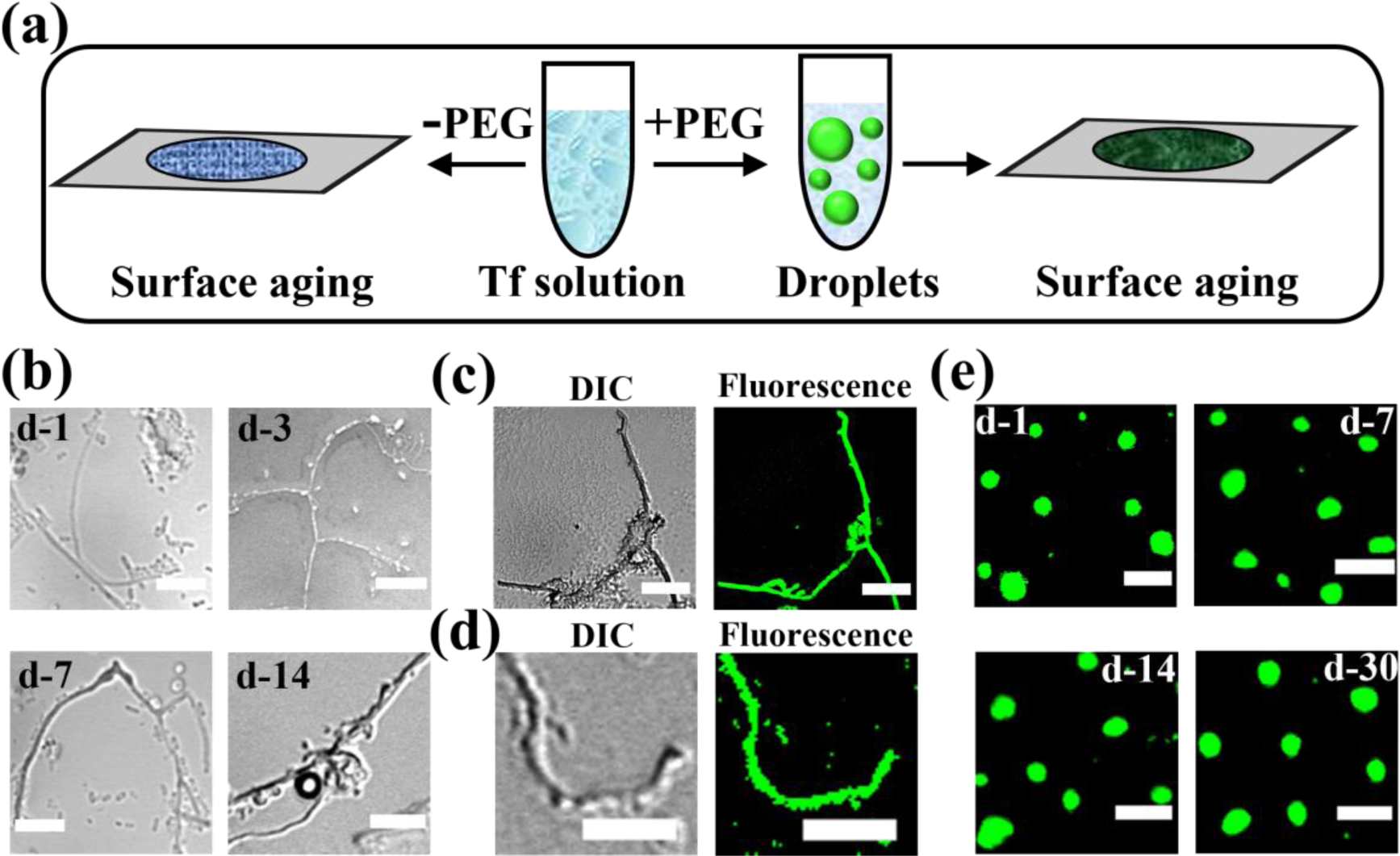
(a) Schematic showing the surface ageing experiments of bare Tf and Tf droplets on the glass surface. (b) Confocal DIC images showing the effect of aging on the surface deposited bare Tf. Confocal (DIC and fluorescence) images of (c) surface deposited FITC-labeled bare Tf on the glass surface and (d) Th-T equilibrated bare Tf. (e) Confocal images of FITC-labelled Tf droplets in the presence of 10% PEG as a function of surface aging. The scale bars correspond to 5 µm.

It is evident that the green emission appears exclusively from the fibrillar aggregates, signifying the presence of Tf inside these aggregates. In order to further establish that these aggregates of Tf are amyloid-like fibrillar aggregates, we performed thioflavin T (Th-T) binding assay with the surface deposited dried Tf samples. Th-T is a well-known amyloid fibril marker and exhibit enhanced fluorescence upon binding with amyloid-like protein aggregates.^68, 69^ We equilibrated 20 µM Th-T with the 1µM aqueous Tf solution for 1 h and subsequently drop-casted on the glass surface. The dried Th-T equilibrated sample was imaged under the CLSM (Figure 5d). It has been observed that Th-T specifically binds with the fibrillar aggregates of Tf and exhibits intense fluorescence upon excitation at 488 nm. Importantly, no such fibrillar aggregates of Tf has been observed inside the aqueous chamber under the same experimental conditions (Figure S13), suggesting that solid-liquid interface has subtle effect on the aggregation of Tf. Therefore, the spontaneous fibrillation of Tf on the glass surface is possibly due to the dehydration and subsequent deformation in its native secondary structure upon surface drying. Notably, 1 µM apo-Tf also exhibits similar fibrillar deposits on the glass surface (Figure S14). Previously, the aggregation and fibrillation process of Tf has been studied on a variety of solid surfaces using a range of experimental techniques.^70–72^ It has been highlighted that solid surfaces play crucial role in fibre formation via deformation of the native structure of the flexible Tf molecules. In addition, Sadler and coworkers have demonstrated that dimer formation promotes fibrillation of Tf on surfaces.^72^ Moreover, it is well-documented in the literature that the extent of protein hydration and the nature of solid supports dictate the aggregation pathways of a wide range of biomolecules.^73–78^

Next, we asked whether the phase separated Tf molecules inside the liquid-like droplets also undergo aggregation and fibrillation upon surface deposition or not. This is particularly a relevant question as previous studies have highlighted the crucial role of liquid-like droplets as active intermediates during protein fibrillations.^17, 18, 21, 27, 28, 32, 38^ It has also shown that the nucleation of protein fibrillation starts within individual droplets. To know the role of the liquid-like condensates of Tf in the fibrillation pathway, we used 10% PEG to initiate phase separation of FITC-labeled Tf in pH 7.4 phosphate buffer at 37 ℃. The phase separated droplets were drop-casted on cleaned cover slipes and incubated for 1-, 7-, 14-, and 30-days at 37 ℃. Subsequently, these dried samples were imaged under the confocal microscope. Interestingly, the confocal image of the d-1 sample shows well dispersed spherical droplets with characteristic green fluorescence with no evidence of any fibrillar aggregates. Similarly, uniform spherical droplets have also been observed with d-7, d-14, and d-30 samples. Notably, the mean size and shape of these droplets remain unaltered even after 30-days of incubation on solid support (Figure S15). Moreover, no liquid-to-solid-like phase transition has been observed upon 30-days of incubation of the aqueous dispersion of these droplets as revealed from the day-light photographs (Figure 5e). The absence of any fibrillar aggregates even with the dried Tf samples along with lack of any liquid-to-solid-like phase transition suggest that these liquid-like droplets act as protective scaffolds for the phase separated Tf from denaturation and subsequent aggregation. Here it should be noted that the droplet interior of biomolecular condensates is highly dynamic and mobile in nature due to the presence of bound water molecules associated with the solvation shell of the native protein.^21, 26, 45^ Our findings indicate that the phase separated Tf droplets can able to retain these bound water molecules even after 30-days of incubation on a solid support without any adverse structural disruptions. This is particularly interesting and unique observation as the protective action of phase separated droplets against dehydration and aggregation of Tf is opposite to what is known in the literature for disease associated IDPs.

To further explore the residue level structural changes within the single droplet as a function of surface aging, we performed single droplet Raman spectroscopy with a laser (*λ* = 532 nm) Raman set-up coupled with a microscope to interrogate individual droplets (Figure S16). Here our aim is to understand the residue level structural changes of fibrils and droplets of Tf upon surface aging for a period of 14-days. Figure 6a compares the individual normalized Raman spectrum recorded for three independent sets (*n* = 3) of Tf fibrils and droplets for d-1 and d-14 surface aged samples. These spectra were corrected from the background signals of 10% PEG (Figure S17). The average spectra for fibrils and droplets are depicted in Figure 6b. It is evident that both fibrils and droplets exhibit characteristic Raman bands for different vibrational modes of the polypeptide chain of Tf. The Raman spectrum of Tf is dominated by amide I (1630–1700 cm^−1^), amide III (1230–1300 cm^−1^), different vibrational modes of aromatic amino acids (tyrosine, tryptophan, and phenylalanine) and aliphatic sidechain vibrations (Figure 6b and Table S1).

**Figure 6.**
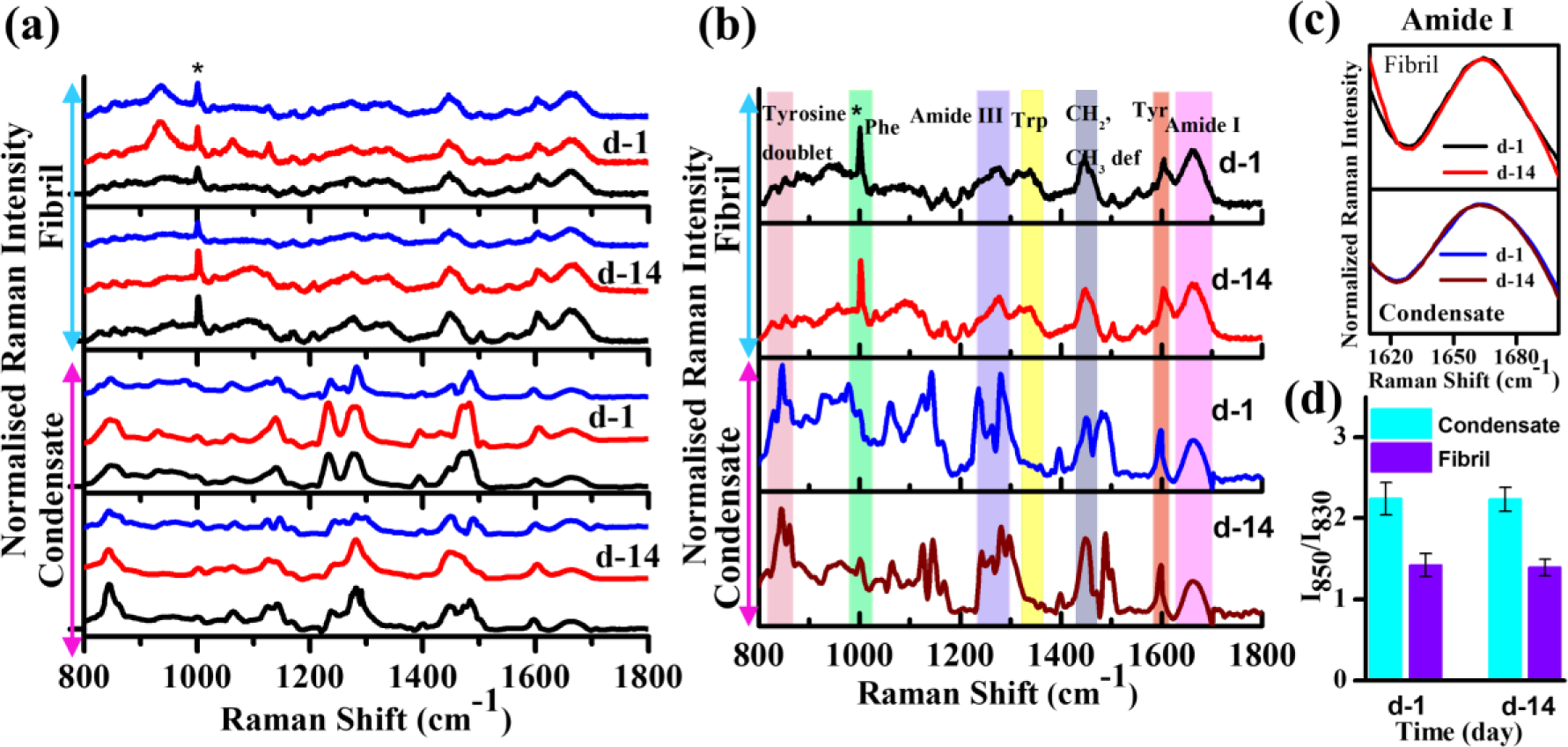
(a) Time-dependent individual (*n* = 3) Raman spectra (*λ* = 532 nm) of Tf fibrils and condensates deposited on the glass slide. Spectra were normalized at 1002 cm^−1^ for comparison. (b) Normalized Raman spectra averaged over three independent runs (*n* = 3) of the Tf fibrils and condensates. Characteristic peaks are marked with colour bars. (c) Normalized amide I spectra of Tf fibrils and condensates at d-1 and d-14. (d) Tyrosine Fermi doublet (I_850_/I_830_) intensity ratios of Tf fibrils and condensates at d-1 and d-14. The data points represent the mean ± s.e.m. for three independent measurements.

The amide I band originates due to the carbonyl (-C=O) stretching and amide III band arises primarily due to the C-N stretching and N-H bending vibrations of the polypeptide chain. These two signature bands in the Raman spectrum are very sensitive to the alteration in the secondary structure of proteins and often used as secondary structure markers.^27, 29, 79^ Notably, the amide I band of Tf fibrils appears at 1662 cm^−1^ and its position does not change significantly upon 14-days of incubation on the glass surface (Figure 6c). Similarly, the full width at half maximum (FWHM) of the amide I band of Tf fibrils shows a value of ∼50 cm^−1^, which remains unaltered upon 14-days of aging. On the other hand, Tf droplets shows a broad amide I band centred at 1664 cm^−1^ with a FWHM of ∼60 cm^−1^ (Figure 6c). Importantly, the position and shape of the amide I band of Tf droplets do not change upon aging for 14-days. The broader amide I band for Tf droplets clearly indicates substantial amount of conformational heterogeneity inside the phase separated droplets of Tf. This argument is further supported by the position and shape of the amide III band. While the amide III band of Tf fibrils appears at 1274 cm^−1^, the same appears at 1284 cm^−1^ for Tf droplets, authenticating considerable amount of conformational heterogeneity inside the phase separated droplets relative to the organized fibrillar assembly of Tf. The intensity ratio of the tyrosine Fermi doublet (I_850_/I_830_) in the Raman vibrational spectrum indicates the hydrogen bonding (H-bonding) propensity between the phenolic hydroxyl moity of tyrosine and the surrounding water molecules and shows a typical value of ≥ 2.0 for a well-hydrated tyrosine residue.^29^ The estimated I_850_/I_830_ ratio for Tf fibrils and droplets is found to be 1.42 and 2.24, respectively (Figure 6d). The ratio is significantly lower than 2.0 for fibrillar aggregates of Tf, suggesting lack of H-bonding interactions due to dehydration and expulsion of interfacial water molecules from the hydrophobic fibrillar assembly of Tf. This argument gains support from the earlier studies on amyloid fibril forming proteins/peptides.^77, 78^ On the other hand, the ratio is greater than 2 for liquid-like droplets of Tf, signifying that the tyrosine residues of phase separated Tf is well-hydrated compared to the more organized fibrillar aggregates.^29^ This is expected as biomolecular condensates are known to contain appreciable amount of solvated water molecules inside their dense phase. More interestingly, this intensity ratio does not decrease significantly for both the fibrils and droplets even upon surface aging for 14-days (Figure 6d). Nevertheless, Raman vibrational modes of Tf highlight the presence of hydrated polypeptide chain with conformational heterogeneity inside the liquid-like droplets and the hydration shells remain intact even after surface aging over a period of 14-days.

The present findings highlight an unknown functional aspect of liquid-like biomolecular condensates of biologically active protein (Scheme 2).

**Scheme 2.**
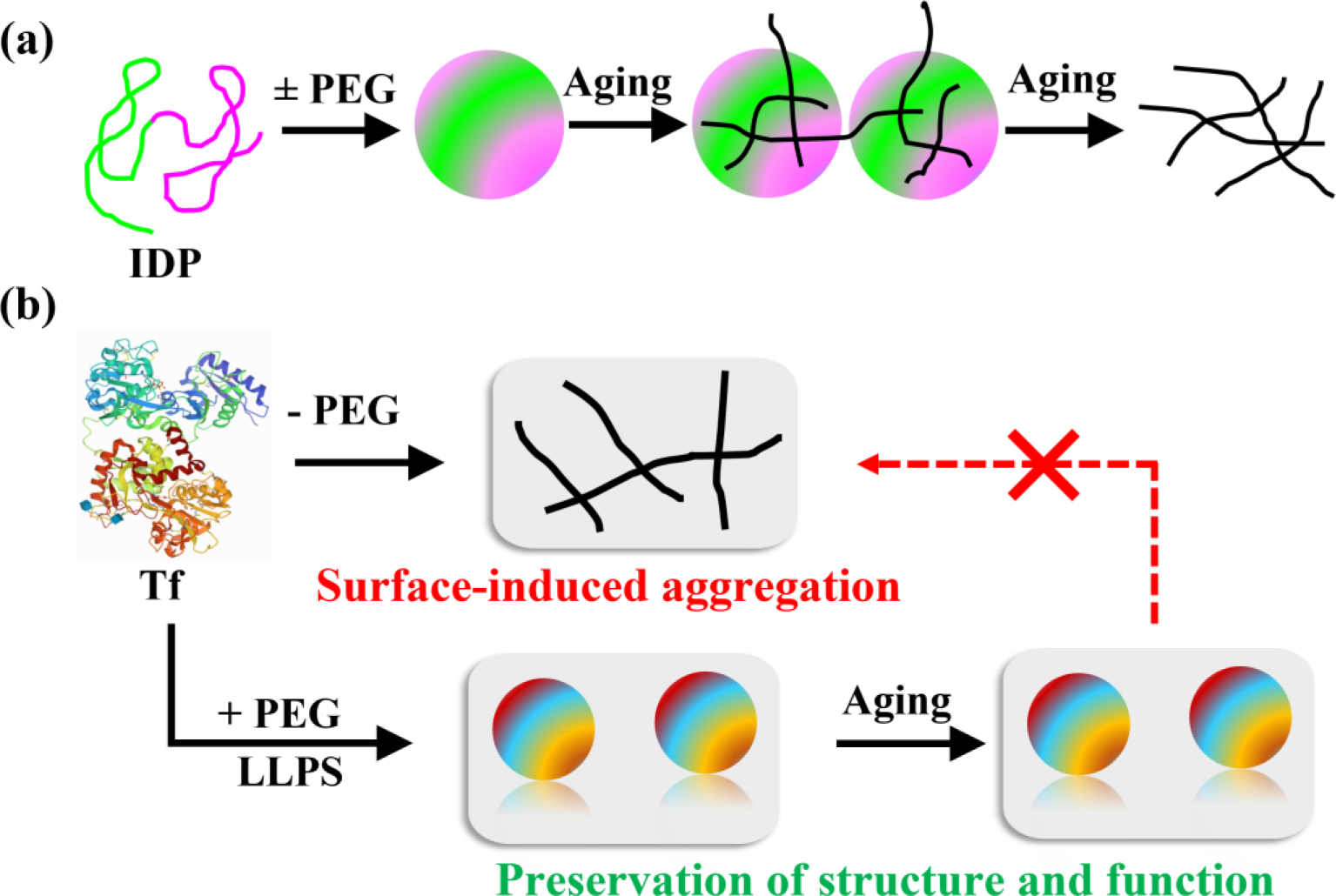
(a) Schematic Illustration of Droplet-Mediated Fibrillation of IDP. (b) Schematic Showing the Effect of PEG as Crowder on the Structure and Function of Tf.

In general, biomolecular condensates of disease associated IDPs containing LCDs are known to undergo fibrillation via liquid-to-solid-like phase transition upon aging (Scheme 2a).

In contrast, for the first time we have demonstrated that biomolecular condensates of Tf act as protective scaffolds to prevent surface-induced pathological fibrillation and the integrity of the phase separated Tf remains intact even after 30-days of aging on the glass surface (Scheme 2b). The liquid-like nature of these condensates with plenty of bound water prevent surface-induced denaturation of phase separated Tf. On the other hand, bare Tf undergoes surface-induced spontaneous fibrillation owing to the deformation of their fragile native conformation as a consequence of dehydration and expulsion of interfacial water molecules which facilitates the formation of hydrophobic fibrillar assembly (Scheme 2b). Although it is a well-known fact that inert crowders are effective stabilizers of the native structure of a wide range of proteins,^80–83^ our present study for the first time discovered that inert crowders stabilize functional protein via inducing LLPS. The present findings are particularly important as protein aggregation and inactivation are major limiting factors for their analytical applications in biotechnology related fields.

## Summary

We have discovered a unique salt-mediated reentrant-type LLPS profile of an ordered functional protein namely, Tf under macromolecular crowding. Both holo- and apo-Tf undergo UCST-based homotypic LLPS via enthalpically driven hydrophobic interactions possibly through the involvement of their 18-residue long LCD containing hydrophobic amino acid residues in the N-lobe. The phase separation of Tf in both the low- and high-salt regimes is solely driven by hydrophobic intermolecular interactions. The secondary structure of Tf does not change significantly upon phase separation even after 14-days of incubation at 37 ℃. The phase separated biomolecular condensates of Tf do not undergo liquid-to-solid-like phase transition over a period of 30-days which is well beyond the cellular half-life (7–8-days) of Tf. More importantly, we have demonstrated an unknown functional aspect of these robust biomolecular condensates of Tf on solid supports. While the bare Tf undergo spontaneous aggregation and fibrillation on the glass surface, the phase separated droplets of Tf completely inhibit the fibrillation process even after 30-days of aging on the glass surface. Owing to the fragile nature, bare Tf suffers conformational deformation upon surface deposition which triggers their aggregation and subsequent fibrillation on the glass surface. On the other hand, structural insights from single droplet Raman spectroscopy reveal conformational heterogeneity inside the phase separated droplets of Tf and retention of bound solvated water inside the membraneless scaffolds even after complete surface drying. The present study highlights the protective role of phase separated droplets toward surface-induced aggregation and fibrillation of Tf which is very unusual considering that biomolecular condensates are known to promote protein aggregation and fibrillation of disease associated proteins/peptides. We strongly believe that our findings may have significant implications in biomedical and biotechnology related fields.

## ASSOCIATED CONTENT

### Supporting Information

The Supporting Information is available free of charge on the ACS website.

Experimental methods and sample preparations; time-lapse DIC images of Tf droplets; confocal image of FITC-labeled apo-Tf droplets; confocal images of FITC-labeled Tf droplets as a function of liquid-phase aging and their corresponding size-distribution histogram; confocal images of FITC-labeled Tf droplets as a function of Tf concentrations; CD spectra of Tf at different pH; confocal images of FITC-labeled Tf droplets as a function of NaSCN, 1,6-hexanediol, (NH_4_)_2_SO_4_, and NaCl concentrations;FTIR spectra of Tf in the absence and presence of different crowders; FESEM image of Tf fibrils; confocal images of FITC-labeled Tf in liquid chamber; confocal images of FITC-labeled apo-Tf fibrils upon 14-days of aging; plot of mean size of Tf droplets as a function of surface aging; photograph of Tf droplets under laser Raman microscope setup; corrected single droplet Raman spectra of Tf droplet; and list of characteristic Ramn peaks of Tf (PDF)

## Notes

The authors declare no competing financial interests.

## Acknowledgment

The authors acknowledge Indian Institute of Technology (IIT) Indore for providing financial support, infrastructure, and instrumentation facilities. The author thanks SIC, IIT Indore, for instrumental facilities. The Raman facility received from the Department of Science & Technology (DST), Government of India, under FIST scheme (Grant no. SR/FST/PSI-225/2016) is highly acknowledged. C.K.P. and C.R. acknowledge the Ministry of Education (MoE) and DST (DST/INSPIRE/03/2019/002160/IF190314) for providing fellowships.

## Notes

### Competing Interest Statement

The authors have declared no competing interest.

