## Supplemental file for "Macromolecular Crowding Promotes Reentrant Liquid-Liquid Phase Separation of Human Serum Transferrin and Prevents Surface-Induced Fibrillation"

### Table of Contents


| S. No. | | Page No. |
|---|---|---|
| 1 | Methods | S3-S5 |
| 2 | Supporting Figures | S6-S22 |
| 3 | Table | S23 |
| 4 | References | S24 |

#### 1. Methods

##### 1.1. Prediction of LCDs and IDRs of Tf

To predict the presence of low complexity domains (LCDs) and disorder regions (IDRs) in amino acid sequence of Tf, we used Simple Molecular Architecture Research Tool (SMART) and IUPred2 respectively.[1,2] The IUPed2 data were plotted using the origin 8.1 software.

##### 1.2. Preparation of Buffer Solutions and Crowder Solutions

All the buffer solutions were prepared in the presence of 0.02% sodium azide and sterilized by autoclaving to minimize the growth of bacteria. Different buffer solutions with pH values of 3.0, 4.0, 7.4, 9.0, and 10.0 were prepared by using Milli-Q water. The buffer strength was kept constant at 50 mM. Sodium citrate buffer (pH 3.0), sodium acetate buffer (pH 4.0), phosphate buffer (pH 7.4), tris buffer (pH 9.0), and carbonate-bicarbonate buffer (pH 10.0) were used individually.

Solutions of different crowders such as 10% (w/v) PEG 8000, 10% (w/v) dextran 70, 12.5% (w/v) Ficoll 400 were prepared from stock solution of 40% (w/v) PEG 8000, 40% (w/v) dextran 70, and 40% (w/v) Ficoll 400, respectively. 20 mg/mL BSA was prepared from the stock solution of 332 mg/mL BSA.

##### 1.3. Labeling of Protein and Crowders with Fluorescent Dyes

The concentration of Tf and apo-Tf was estimated spectrophotometrically at $\lambda$ = 280 nm using the reported extinction coefficient of $1.04 \times 10^5$ and $8.50 \times 10^4$ $M^{-1}$ $cm^{-1}$, respectively.[3] Tf was labeled with FITC dye according to an earlier reported method.[4] In short, 1 µM Tf was mixed with FITC in a molar ratio of 1:10 ([Tf]: [FITC]). The mixture was incubated for 4 h at room temperature followed by 6 h at 4 °C on a magnetic stirrer with constant speed (250 rpm). After the completion of the reaction, the unconjugated dyes were removed using dialysis (molecular weight cut-off 3.5 kDa) against 50 mM PBS at 4 °C for 12 h with regular buffer exchange in 2 h interval. The same procedure was followed for the labeling of PEG and BSA with RBITC.

##### 1.4. Sample Preparation

###### Liquid Phase Aging

Bare Tf and Tf droplet solutions were kept at 37 °C inside an incubation chamber for required days of aging. For CLSM, 20 μL aliquot was withdrawn from the solutions and drop-cast over cleaned glass slide and then immediately covered with cleaned coverslip. The coverslips and glass slides were first cleaned with 2% Hellmanex III and then with chromic acid. Each of these cleaning steps was followed by repeated washing with Milli-Q water. Finally, these washed slides and coverslips were rinsed with methanol and dried in vacuum oven. The edges were sealed with nail paint to avoid evaporation. Here it is important to mention that small amount of nail paint at the edges does not interfere with the liquid-liquid phase separation (LLPS) of Tf.

###### Surface Aging

For surface aging experiments, 20 μL aliquot of Tf fibril (1 μM Tf in phosphate buffer) and Tf droplet (1 μM Tf + 10% PEG in phosphate buffer) were drop-cast over a cleaned glass slide and kept inside the incubation chamber at 37 °C for 1- to 30-days.

##### 1.5. LLPS Assay of Tf in the Presence of Crowders

LLPS of Tf was examined in the presence of different crowders (PEG 8000, dextran 70, Ficoll 400, and 20 mg/mL BSA) as a function of Tf concentration (5–50 μM), crowder concentration (1–20%), and incubation time (1–14 days) at 37 °C. These samples were prepared in 5 mL glass vials and then kept at 37 °C in an incubation chamber.

The salt-dependent study was performed by adding different concentrations of NaCl (0-3 M), NaSCN (0-2 M), and 1,6-hexanediol (0-10%) from their respective stock solutions.

The individual solutions were prepared by adding different concentrations of salts/aliphatic alcohol in aqueous mixture (pH 7.4 phosphate buffer) of 1 µM Tf and 10% PEG 8000. Next, the solutions were kept at 37 °C for 1-day and subsequently measurements were performed.

For the temperature-dependent study, samples (1 µM Tf in the presence of 10% PEG 8000) were prepared in pH 7.4 phosphate buffer at different temperature of 4, 25, 37, and 50 °C and incubated for 1-day. Similarly, pH-dependent study was carried out using 1 µM Tf in the presence of 10% PEG 8000 in different buffers with pH of 3.0, 4.0, 7.4, 9.0, and 10.0 at 37 °C.

**1.6. Th-T Binding Assay**

Stock solution of 2 mM Th-T was prepared in pH 7.4 phosphate buffer. 20 µM Th-T was equilibrated with 1µM aqueous Tf solution for 1 h and subsequently, 20 µL aliquot was drop-cast onto the cleaned coverslip. The dried Th-T equilibrated sample was imaged under the CLSM with an excitation wavelength of 488 nm. The emission was collected in the wavelength range of 490–550 nm.

### Supporting Information Figures

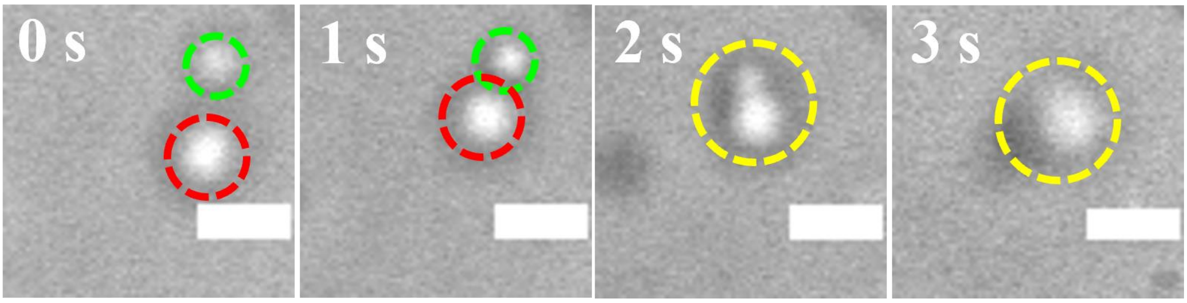


**Figure S1.** Time-lapse confocal (DIC) images of droplets showing a fusion event over a period of 3s. The scale bars correspond to 2 μm.

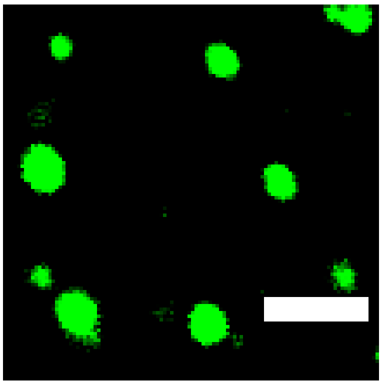

**Figure S2.** Confocal image of FITC-labeled 1 μM apo-Tf in the presence of 10% PEG incubated in pH 7.4 phosphate buffer solution at 37 °C for 1-day. The scale bar corresponds to 5 μm.

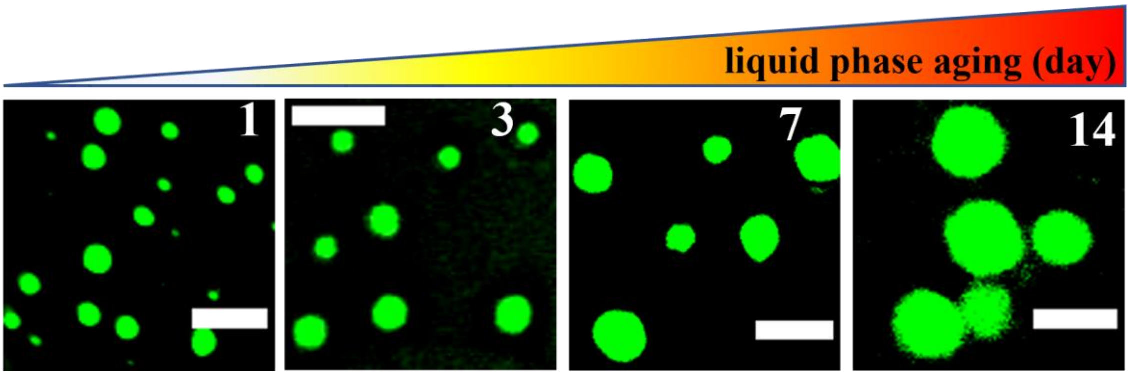


**Figure S3.** Time-dependent confocal images of FITC-labeled Tf (1 μM) droplets in the presence of 10% PEG incubated in pH 7.4 phosphate buffer solution at 37 °C. The scale bars correspond to 5 μm.

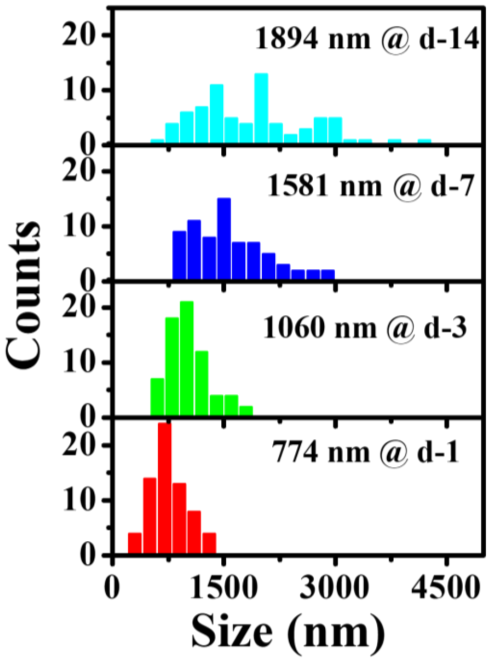


**Figure S4.** Size distribution histograms showing the mean size of the Tf (1 μM) droplets in the presence of 10% PEG under liquid phase aging.

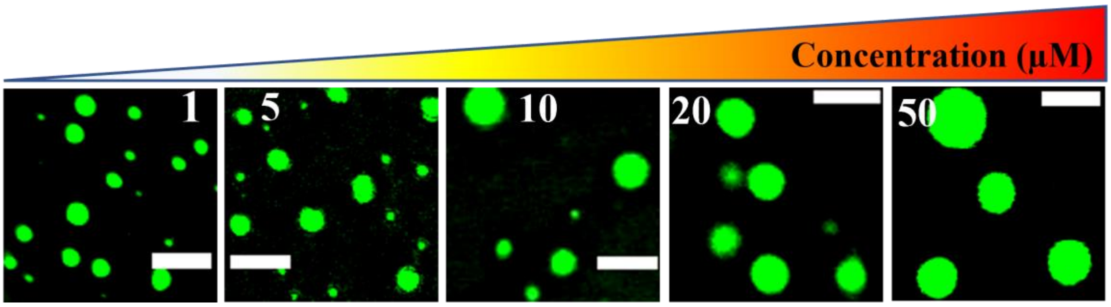


**Figure S5.** Concentration-dependent confocal images of FITC-labeled Tf (1, 5, 10, 20, and 50 µM) droplets in the presence of 10% PEG. The scale bars correspond to 5 µm.

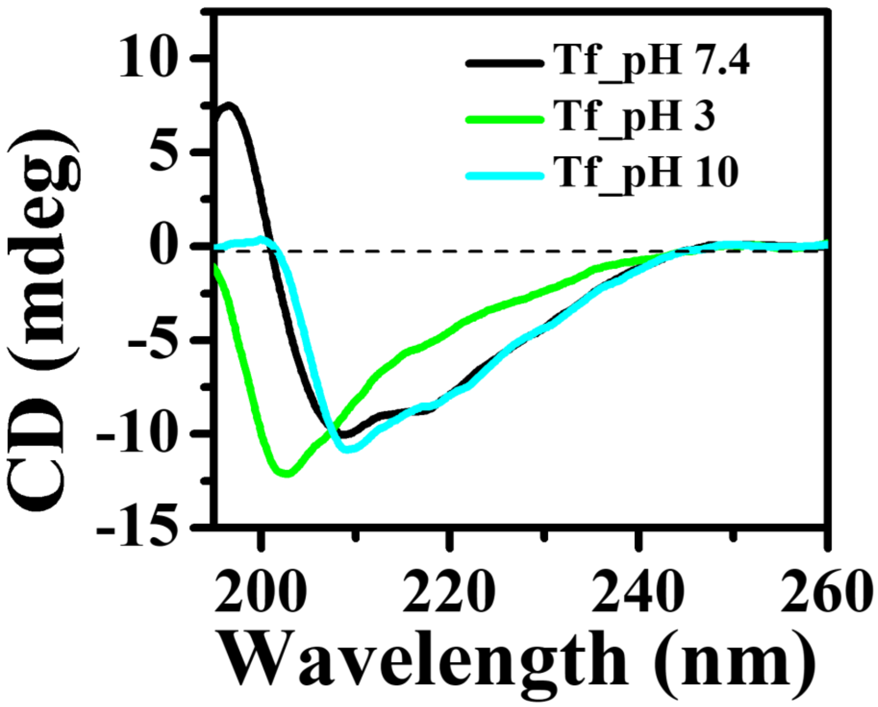


**Figure S6.** CD spectra of 1 μM Tf at different pH upon 1-day of liquid phase aging.

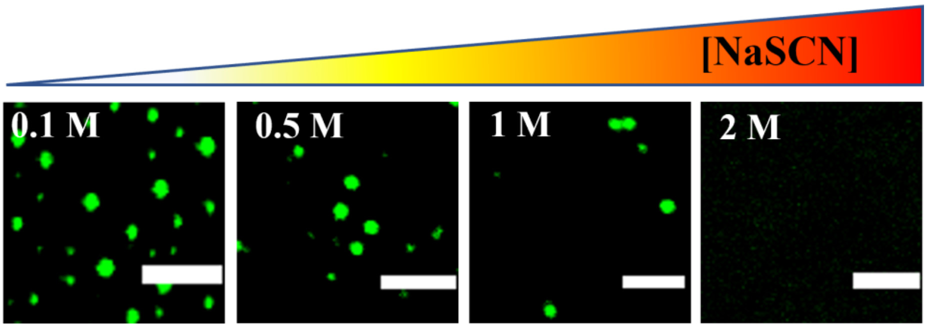


**Figure S7.** Confocal images of FITC-labeled Tf (1 μM) droplets in the presence of 10% PEG as a function of NaSCN concentrations at 37 °C. The scale bars correspond to 5 μm.

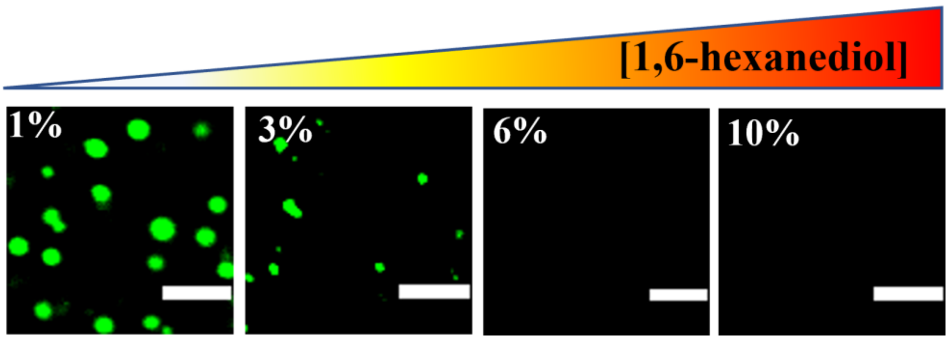


**Figure S8.** Confocal images of FITC-labeled Tf (1 μM) droplets in the presence of 10% PEG as a function of 1,6-hexanediol concentrations at 37 °C. The scale bars correspond to 5 μm.

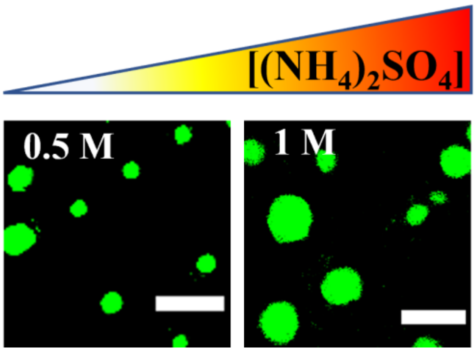


**Figure S9.** Confocal images of FITC-labeled Tf (1 µM) droplets in the presence of 10% PEG as a function of $(NH_4)_2SO_4$ concentrations at 37 °C. The scale bars correspond to 5 µm.

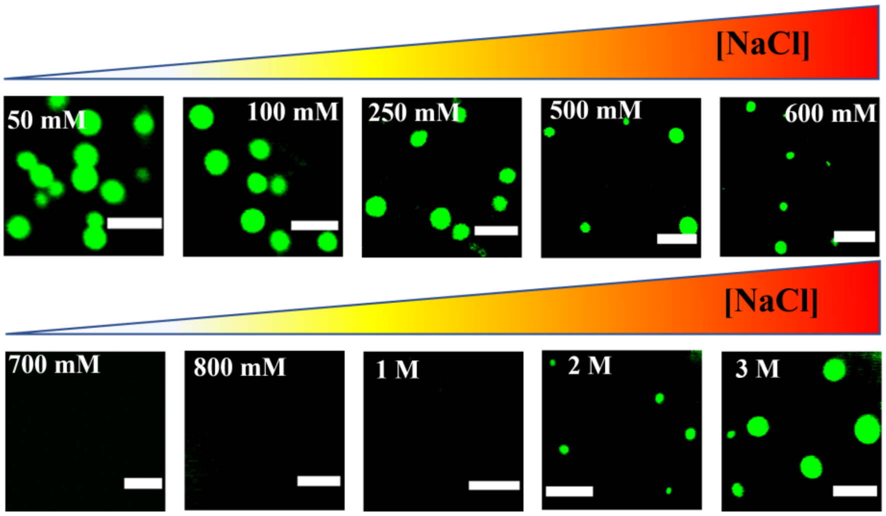


**Figure S10.** Confocal images of FITC-labeled Tf (1 µM) droplets in the presence of 10% PEG as a function of NaCl concentrations at 37 °C. The scale bars correspond to 5 µm.

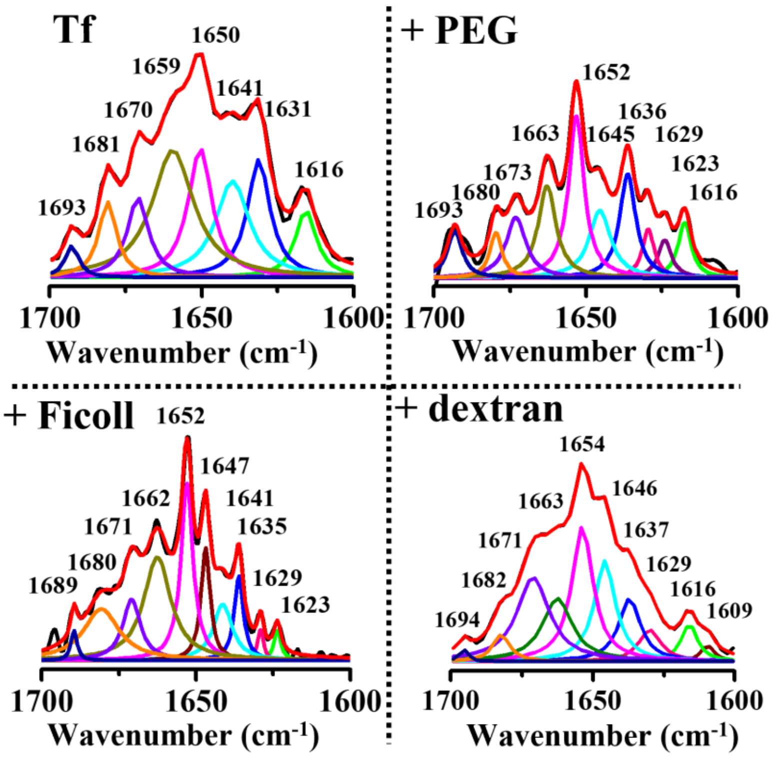


**Figure S11.** FTIR spectra of 50 μM Tf in the absence and presence of different crowders upon aging (1-day) in liquid phase.

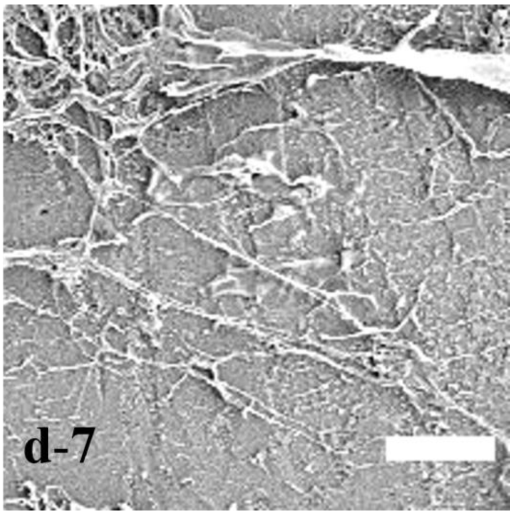


**Figure S12.** FESEM image of 1 µM Tf in the absence of crowders showing fibrillar aggregates after 7-days of aging. The scale bars correspond to 5 µm.

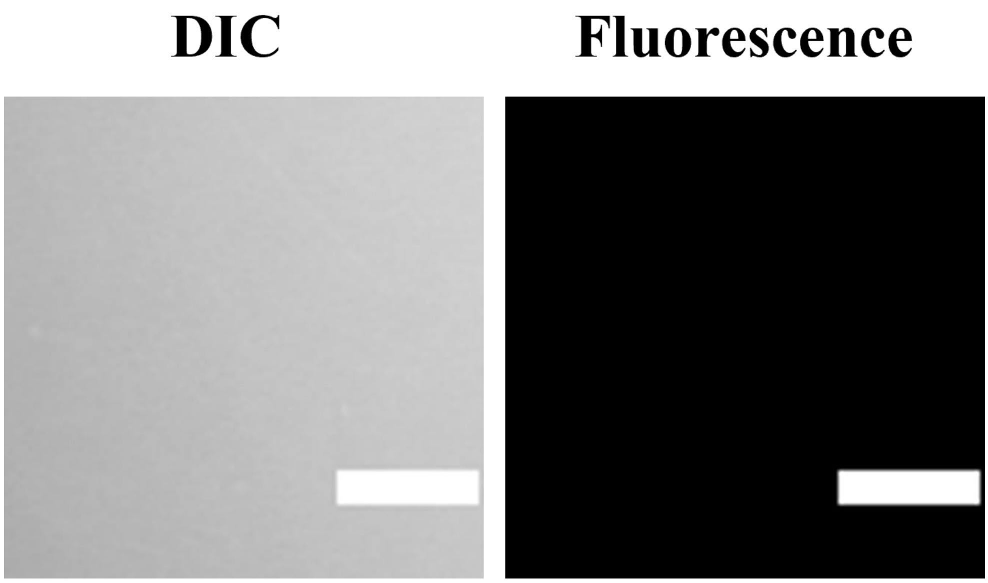


**Figure S13.** Confocal (DIC and fluorescence) images of FITC-labeled 1 µM Tf in the presence of 20 µM Th-T showing no fibrillar aggregates in liquid chamber upon 488 nm excitation. The scale bars correspond to 5 µm.

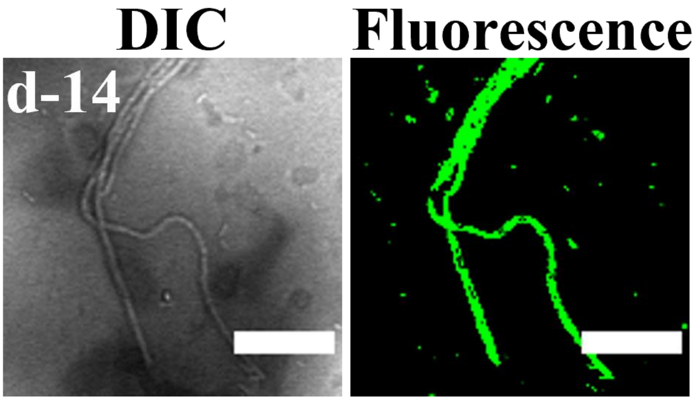


**Figure S14.** Confocal images of FITC-labeled 1 μM bare apo-Tf showing fibrillar aggregates after deposition on the glass surface and surface aged for 14-days. The scale bars correspond to 5 μm.

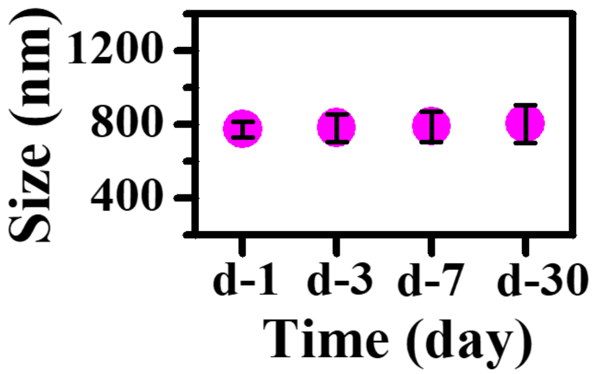


**Figure S15.** Variation in the mean size of Tf droplets as a function of surface aging.

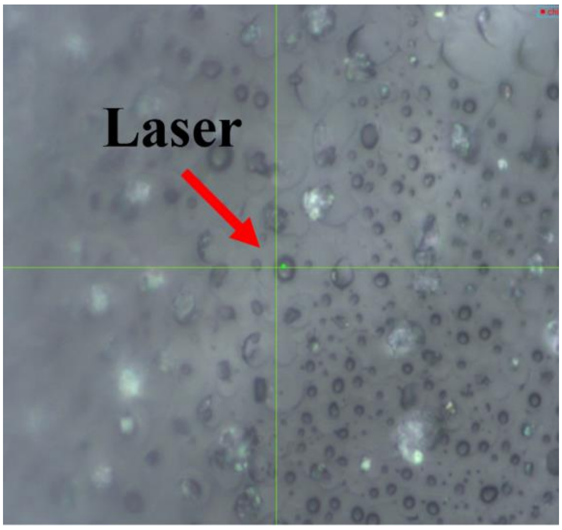


**Figure S16.** Photograph showing the focal point of the laser within a single Tf (50 µM) droplet visualized through the Raman microscope.

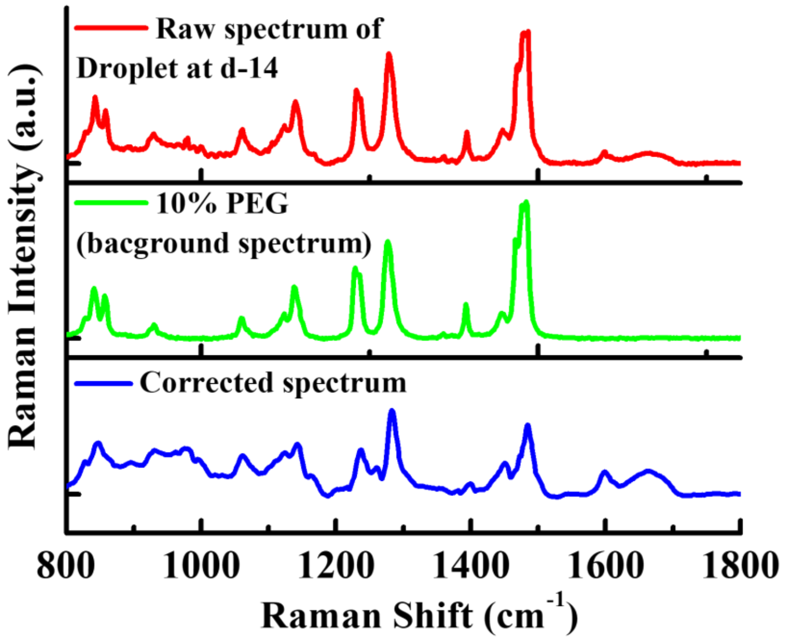


**Figure S17.** Corrected **s**ingle droplet Raman spectra of Tf droplet in the presence of 10% PEG. The upper panel shows the raw spectrum of Tf droplet at day 14, middle panel shows the background spectrum of 10% PEG, and bottom panel shows the background corrected spectrum of Tf droplet.

**Table S1. Characteristic Raman Peaks of Tf**

| Raman modes | Raman peaks |
|---|---|
| **Amide I** | |
| **a. Fibrils** | 1662 $cm^{-1}$ |
| **b. Condensate** | 1664 $cm^{-1}$ |
| **Amide III** | |
| **a. Fibrils** | 1274 $cm^{-1}$ |
| **b. Condensate** | 1284 $cm^{-1}$ |
| **Phenylalanine** | 1002 $cm^{-1}$ |
| **Tryptophan** | 1336 $cm^{-1}$ |
| **Fermi doublet** | 1360 and 1340 $cm^{-1}$ |
| **Tyrosine** | |
| **a. Fermi doublet** | 850 and 830 $cm^{-1}$ |
| **b. Ring stretching mode** | 1600 $cm^{-1}$ |
| **Backbone $CH_2$/$CH_3$ deformations** | 1449 $cm^{-1}$ |
